# Simple and efficient auditory-nerve spike generation

**DOI:** 10.1101/2023.05.02.539135

**Authors:** Alain de Cheveigné

## Abstract

This brief paper documents a simple and efficient method to generate auditory-nerve spike trains for the purpose of simulating neural processes of auditory perception. In response to sound, each auditory nerve fiber carries information to the auditory brainstem in the form of a train of spikes (action potentials), the timing and rate of which reflect the sound. The generation process is usually approximated as Poisson process with a time-varying rate, further modified by refractory effects. The purpose, here, is to simulate spike generation as a time- and interval-dependent thinning process applied to a homogenous Poisson process, allowing for fast generation, cheap storage, and unlimited temporal resolution.

## Introduction

The ear serves to transduce acoustic vibrations in the air to neural patterns and deliver them to the brain, ultimately yielding a sound-evoked percept or behavior. Sound vibrations are transmitted via the outer and middle ear to the inner ear, where they undergo active amplification and filtering, the outcome of which determines the probability of release of quanta of neurotransmitter at the synapse between inner hair cell and auditory nerve fiber, thus determining the probability of occurrence of a spike within that fiber. Information available to the brain about sound is carried collectively by ∼30000 fibers (∼10 fibers contact each of ∼3000 inner hair cells), each carrying a spike train of up to ∼300 spikes/s average rate.

The spike-generation process has been described as an *inhomogenous Poisson process* with a time-varying rate that follows the deflection (and/or velocity) of the basilar membrane at the position of the hair cell, further modified by *refractory effects* that depress firing probability immediately after each spike. This widely used approximation is reviewed by Delgutte (1996), some of its limits are discussed by Heil and Peterson (2017).

Auditory models (or model suites) have been developed to approximate auditory transduction (Osses et al 2022). They serve two purposes: to embody knowledge accrued about the peripheral auditory system (and test it), and to generate patterns of auditory-nerve discharge for simulation of downstream processes within the brainstem. A typical auditory model might combine a linear acoustic filtering module to represent the outer and middle ear, a non-linear filter-bank module to represent cochlear filtering, an inner hair-cell module to represent transduction from vibration to intracellular potential, and a *spike generation* module that generates an array of spike trains.

Simpler configurations may be used for specific tasks, for example by omitting the initial filtering, or reducing the number of channels, or replacing non-linear by linear filters. While a complete, physiologically-realistic model is a worthy goal, a simpler model is often sufficient, or even preferable because more constrained (fewer parameters) and computationally less expensive. In particular, the final spike-generation phase is often omitted.

Nonetheless, generating actual spike trains (as opposed to a time-varying probability) may be useful to assess the impact of refractory effects (not fully captured by the driving probability), or to drive models that rely on processing the spikes themselves. These include coincidence-based models for example for pitch or binaural unmasking, as well as a wider class of spike- or event-based models of perception or information processing (e.g. Liu et al 2019). Spike generation enables a model to simulate the effects of *deafferentation* (Kujawa and Liberman 2009), putatively a major component of hearing loss, for example in terms of stochastic under-sampling (Lopez-Poveda 2014).

Two obstacles to spike generation are computational cost and storage cost, both linked to the common time-sampled simulation approach (e.g. one sample every 10μs). According to the inhomogenous Poisson model with refractory effects, spike probability at each time is determined by the product of a time-varying driving function, and a recovery function dependent on time since the latest spike. This can be simulated by drawing a random number at each time step and comparing it with that probability. Downsides are (a) the large number of random numbers required if the sampling rate is high, (b) the space required to store the (very sparse) time series of values (1 for a spike, 0 otherwise) and (c) the limited time resolution of each spike, dependent on the sampling rate.

Two approaches are available to more efficiently simulate an inhomogenous Poisson process. Both rely on the fact that intervals between events of a *homogenous* Poisson process are distributed exponentially. The two approaches are *time transformation* (e.g. Jackson et al 2005) and *thinning* (e.g. Lewis and Shedler 1979). With time transformation, the duration of each randomly-drawn inter-event interval is scaled by a factor that depends on the instantaneous probability at (presumably) the first event of that interval. With thinning, the *maximum* value of the time-varying rate is used to simulate a homogenous Poisson process, and the resulting event train is then *thinned probabilistically* depending on the instantaneous rate at each event of this train.

The time-transformation approach might seem preferable because it requires a single random draw per final spike, whereas thinning is more wasteful. However, time transformation is subject to a subtle issue: the underlying time-varying rate is *sparsely* sampled by the event train, all the more so as the rate is *low*. For example, if an event occurs at a dip in the time-varying rate, the next sampling point will be distant in time, possibly missing an intervening section of high rate. Thinning does not have this problem (at least, not to the same degree), and is the approach used here.

## The toolbox

The toolbox, implemented in Matlab, contains a small set of routines to create and analyze spike trains. A spike train is represented as an array of floating-point numbers, each number representing time relative to a time reference locked to the beginning of the train.

Spike trains are generated according to the thinning method. The input time series (instantaneous rate aka driving function) is scanned for its maximum value, this value parametrizes a homogenous Poisson process that produces a train of inter-spike intervals at that maximum rate. These are cumulatively summed (assuming a first spike at time 0) to obtain a train of spike times. This spike train is then thinned probabilistically depending on the value of the nominal instantaneous rate at the time of each spike, to simulate the time-varying Poisson process. Finally, the spike train is again thinned probabilistically according to a recovery function. Initial spike generation and thinning are vectorized and thus fast, but the final thinning requires a loop.

The toolbox includes a few additional routines to modify the spike train (jitter, cancellation filter, c.f. de Cheveigné 2021, 2023), and for routine statistics such as first-order and all-order inter-spike interval histograms, peri-stimulus time (PST) histograms, and cross-coincidence histograms).

The toolbox is mainly intended as an add-on to existing auditory model toolboxes (e.g. Osses et al. 2022).

The toolbox is archived at https://zenodo.org/badge/latestdoi/635354073.

## Results

### Spike generation

Figure 1 illustrates the generation process. A homogenous Poisson process with a rate equal to the peak rate of the driving function (top) produces a train of spikes (black dots), that is thinned, first on the basis of the instantaneous values of the driving function (blue), and next on the basis of the recovery function (red).

**Fig. 1.**
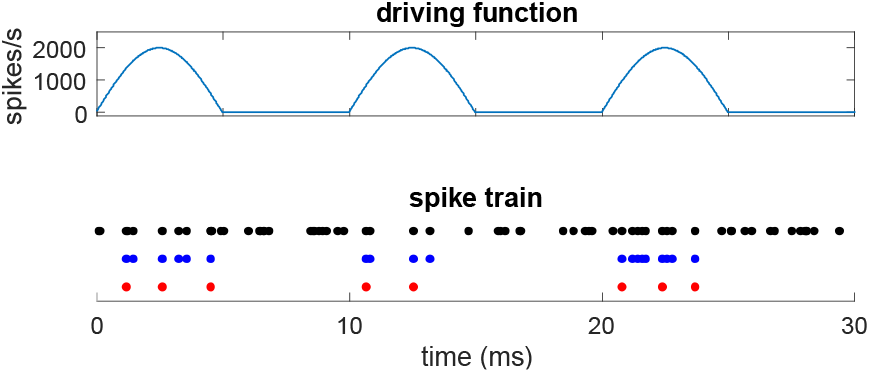
Top: driving function (half-wave rectified 100 Hz sinusoid). Bottom: spike train produced by a homogenous Poisson process of rate 2000 spikes/second (black), the same after thinning according to the driving function (blue), and after additional thinning according to the recovery function (red).

### Constant driving function

In the absence of stimulation, most auditory nerve fibers have a spontaneous activity that can be characterized by a driving function that is constant (at least approximately, c.f. Heil and Peterson 2017). Stationary stimulation at a high frequency, beyond the phase-locking limit (approximately 5 kHz) also entails a constant driving function. Figure 2 shows interval statistics for constant drive, in the absence (black) or presence (blue and red) of refractory effects. The driving rate was 400/s, the effective rate (after thinning) was ∼290/s and ∼260/s for the stepwise (blue) and piecewise linear recovery time (red) respectively.

**Fig. 2.**
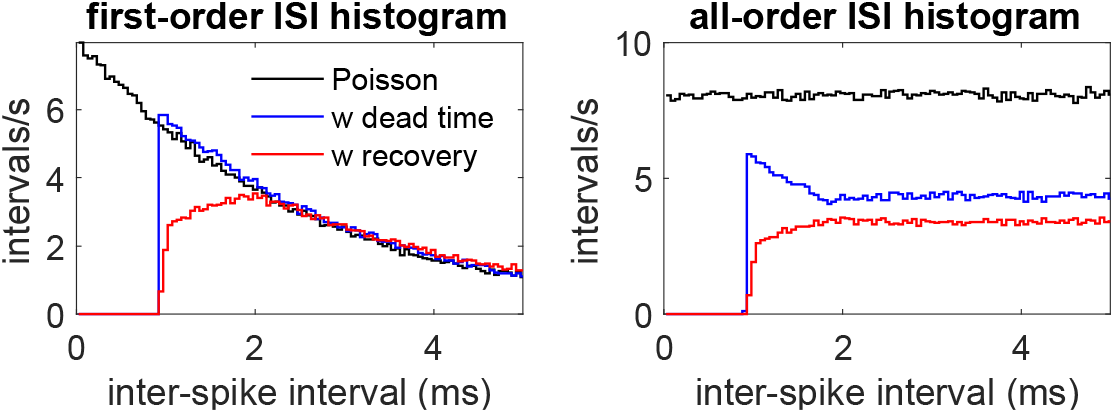
Interval statistics for spike trains resulting from a constant driving function, in the absence (black) and presence (blue and red) of refractory effects. Left: first-order interval histogram, right: all-order interval histogram, binwidth is 0.05 ms. For the blue curve (“dead time”), the recovery function was a step function equal to 0 below 0.8 ms and 1 beyond. For the red curve, the recovery function was piecewise linear (0 below 0.8 ms, 0.5 at 1 ms, 0.9 at 2 ms, and 1 at 5 ms and beyond). Statistics were gathered over a duration of 600 s.

### Halfwave-rectified driving function

Figure 3 (left) shows the peristimulus (PST) histogram for a spike train elicited by a half-wave rectified sinusoidal (100 Hz) driving function in the absence (black) or presence (red) of refractory effects (peak rate 500 spikes/s, average spiking rates 160 and 110 spikes/s respectively). The center and right panels show first-order and all order ISI histograms, respectively. Refractory effects (red) reduce the number of spikes and deplete the zero-order mode of the interval histograms. There is also a slight distortion of the shape of the PSTH, explored in more detail in the next paragraph.

**Fig. 3.**
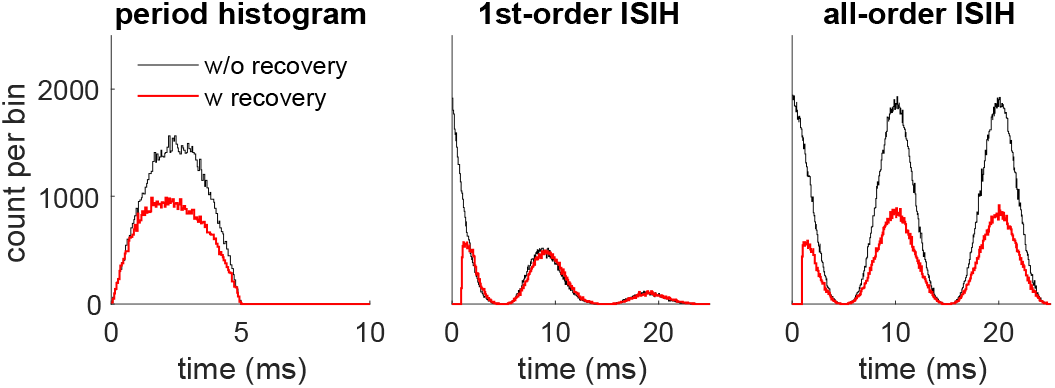
Peri-stimulus time and interval statistics for spike resulting from a half-wave rectified sinusoidal driving function (100 Hz) of peak rate 500 spikes/s, in the absence (black) and presence (red) of refractory effects. The same piecewise-linear recovery function was used as in Fig. 2. Statistics were gathered over a duration of 600 s.

### Side-effects of refractoriness

In the absence of refractory effects (pure non-homogenous Poisson process), the empirical firing rate would faithfully follow the driving function (apart from stochastic variability). The presence of refractory effects induces some distortions that are worth recognizing, so as to avoid attributing them to upstream phenomena (e.g. adaptation).

Figure 4 shows three examples of such effects. The left panel shows a peri-stimulus time histogram (PSTH) in response to 5000 repetitions of a pulse of constant driving rate, either 1000 spikes/s (black) or 5000 spikes/s (red) (effective rates 410 and 730 spikes/s respectively). A small onset overshoot is visible at the lower rate, at the higher rate it takes the form of a ringing pattern with a period close to the absolute refractory period (∼1 ms). The center panel show the all-order ISIH for the same two spike trains, again showing evidence of periodicity at the higher rate. The right-hand panel shows a period histogram in response to a half-wave rectified 200 Hz sinusoidal driving function. The peak driving rate was either 1000 spikes/s (black) or 5000 spikes/s (red) (effective rates 180 and 360 spikes/s respectively). At the lower rate (black) the histogram is a slightly distorted version of the driving function (as in Fig. 3 left), at the higher rate the peak splits into two.

**Fig. 4.**
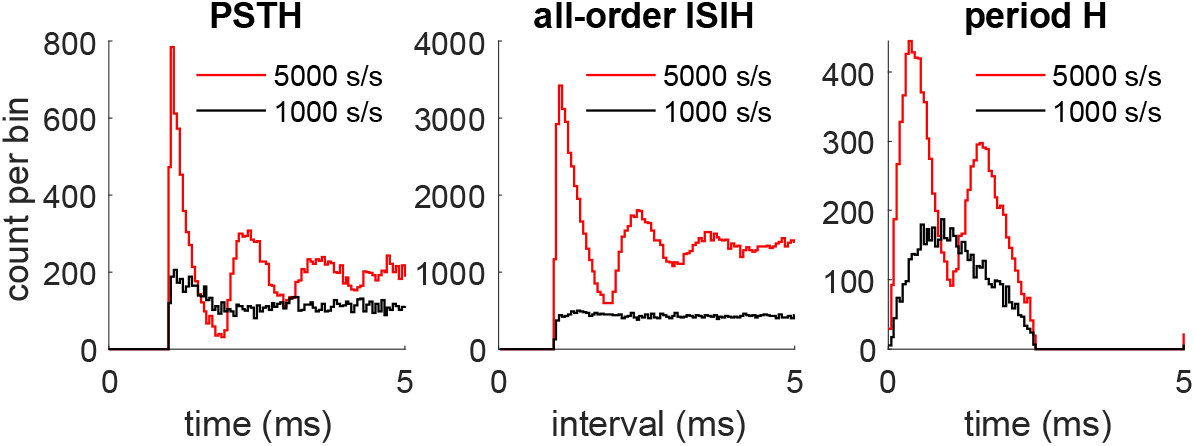
Side-effects of refractoriness. Left: peristimulus time histogram at the onset of a short pulse of constant rate repeated 5000 times, The nominal driving rate was 1000 spikes/s (black) or 5000 spikes/s (red) (effective rates were 410 and 730 spikes/s respectively).

In such conditions, one might be tempted to infer properties of the driving function (reflecting the stimulus, the filter, adaptation processes, etc.) (Lopez-Poveda 2005) that are instead an artifactual side effect of refractoriness. These effects are prominent mostly for high peak driving rates.

### Jitter

Adding jitter to spike times is a convenient way to model loss of synchrony at higher frequencies. Loss of synchrony is usually attributed to low-pass filtering effects due to inner hair-cell capacitance (Russell and Sellick 1983), but its effects are quite well captured by adding a Gaussian jitter to each spike time (Fig. 5) (compare with e.g. Versteegh et al 2011). Adding Gaussian jitter is thus an expedient way to reproduce phase-locking roll-off at higher frequencies, at least for the purpose of driving downstream processing models.

**Fig. 5.**
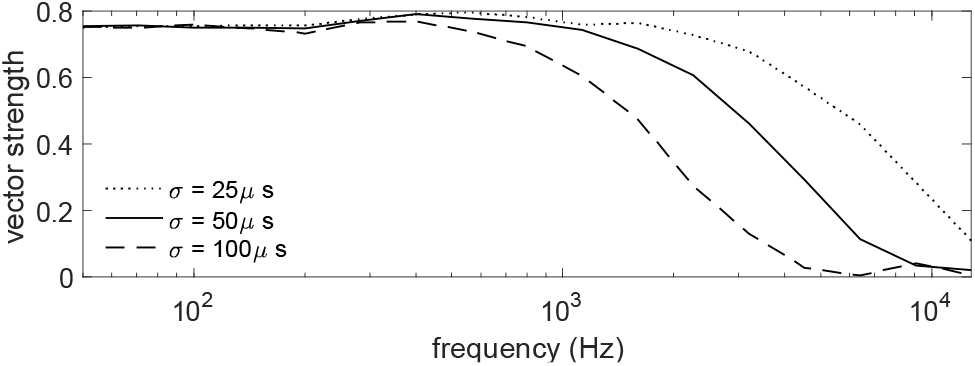
Effect of adding Gaussian jitter to spike times on vector strength, for a half-wave rectified sinusoidal driving function (1000 spikes/s peak rate), for several values of standard deviation. A value of ∼50μs mimicks quite well the synchrony roll-off observed in auditory-nerve recordings in the cat. The subtle increase in vector strength circa ∼400 Hz is a side-effect of refractoriness.

### Cancellation

The cancellation filter involves gating one spike train based on another spike train. Specifically, any input spike that coincides with a gating spike (within a window) is removed. It can be seen as a form of thinning by which spikes are removed to change the temporal structure of the spike train. If the gating spike train is the same as the input but with some delay (or advance), the filter performs *harmonic cancellation* (de Cheveigné 1993, 2021, 2023).

Figure 6 illustrates this process applied to a spike train generated from the half-wave rectified sum of two sinusoids, 80 Hz and 100 Hz respectively. The top plot shows an all-order ISI histogram calculated from the raw spike train. The middle and bottom plots show the result of applying a cancellation filter with delay equal to 10 ms and 12.5 ms respectively (inverses of the two frequencies). The first filter appears to effectively suppress correlates of 100 Hz, and the second those of 80 Hz. Of course, the same effect could be obtained by filtering *before* spike generation (for example cochlear filtering), but it is interesting to see a similar result from processing in the neural domain.

**Fig. 6.**
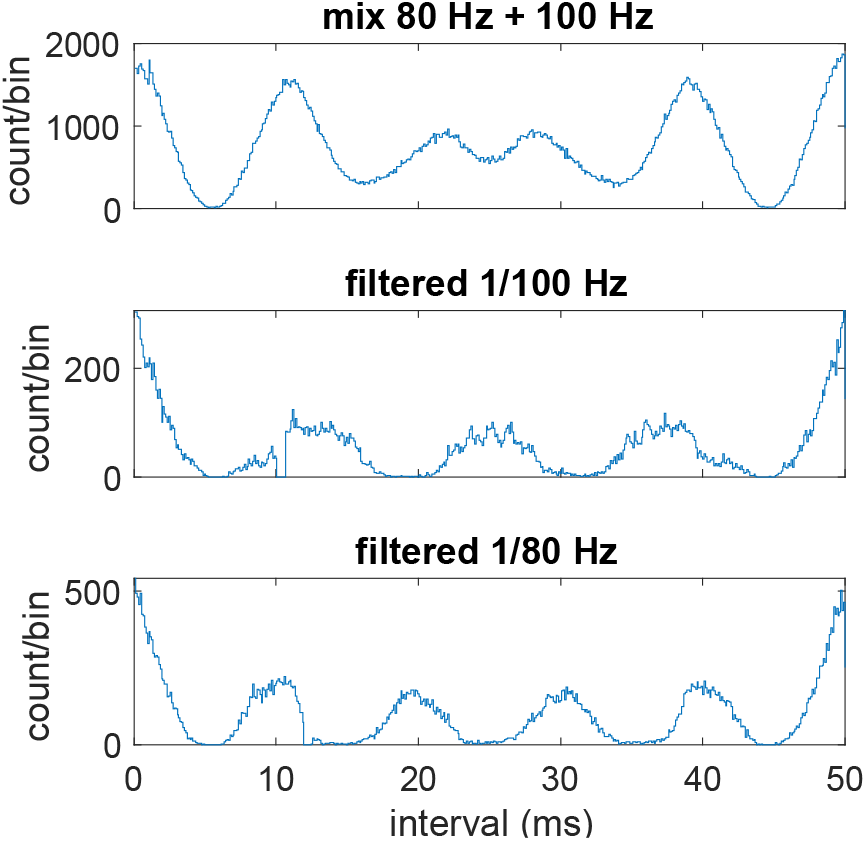
Neural cancellation filter. Top: all-order inter-spike interval histogram for a driving function consisting of the half-wave rectified sum of 80 Hz and 100 Hz sinusoids. Middle and bottom: same, after filtering the spike train with a cancellation filter of delay parameter 10 ms or 12.5 ms (inverses of 80 Hz and 100 Hz).

The peak firing rate here was 1000 spikes/s, a 1 ms step recovery function was applied, and 10 spike trains were generated and merged (compound rate 1940 spikes/s). The cancellation filter kernel was boxcar-shaped with a width of 1 ms.

### Computation and storage

Computation time is data- and parameter-dependent, but to give an idea, simulation of a full set of 30000 fibers takes roughly ten times real time on a 2.4 GHz 8-core MacBook Pro.

At 100 spikes/s average output rate, the resulting spike trains require 24 Mbytes storage (double precision) as opposed to 375 Mbytes if the spike trains were stored as a one-bit time series with 100 kHz sampling rate, or more for an integer or float time series.

### The toolbox

The Matlab toolbox is available at XXX. Most routines follow the same conventions: calling the routine without output arguments produces a plot reflecting the outcome (in some hopefully helpful fashion), calling it without input arguments runs a short piece of example code, hopefully useful to understand the effects of the routine and how to deploy it.

## Summary

This paper documents simple and efficient methods of spike generation and processing for the purpose of modeling peripheral transduction and downstream neural mechanisms. The methods are not new, but they are not often deployed in the context of auditory model simulations, where they can be useful given the high rate and large volumes of neural patterns produced by the roughly 30 000 fibers of the human auditory nerve. Some auditory modeling suites already offer spike train output; a benefit of the present may be to add that capability to those that do not.

The main benefits are *fast generation* (relative to the uniform time-series approach that a decision to fire or not at each time step), *reduced space requirements* (relative to a sparse time series of pulses), and *precise time resolution* (limited by floating-point resolution, rather than sampling rate). Spike times can easily be converted to a time series of pulses, for example for the purpose of autocorrelation or cross-correlation, although the toolbox offers time-based alternatives for those operations.

Time-stamped event-based representations offer theoretical advantages, related to the potentially large amount of information carried by each spike (limited only by the precision of its representation, Maass 2001), and are the focus of intense interest for computational, sensory and cognitive applications (Maass 2015, Liu et al 2019).

*Thinning* is the key to fast generation of arbitrarily-timed events from a Poisson process with time-varying rate and/or refractory effects. Thinning was also used by the cancellation filter to selectively modify the stimulus-driven temporal structure of the spike train (e.g. suppress a periodic masker). As another example, a Poisson-distributed spike train can be transformed by thinning into a Poisson + gamma mixture distribution, which better models spontaneous auditory-nerve interval distributions (Heil et al 2007). Thinning is quite versatile.

*Time-transformation* (Jackson and Carney 2005), in principle yet more efficient than thinning, faces the problem that a momentarily low value of the rate will cause the next spike to be distant in time, potentially missing an intervening increase of the rate function that should normally have resulted in spiking. With thinning, the driving function is sampled “uniformly” (albeit stochastically) at the highest rate. There remains the problem that, after thinning, low-amplitude portions of the driving function are more sparsely sampled than high, but this is inherent in stochastic sampling and unrelated to the simulation method.

Refractory effects may modify the relation between the driving function and the empirically-observed spiking rate, particularly at high instantaneous rates. This modification may affect statistics derived from experimental measurements (Fig. 4), and possibly also properties of downstream neural processing. Phenomena such as fast initial adaptation, ringing, or peak splitting are unlikely to result from refractoriness in general, but it is worth keeping that factor in mind when investigating them. Merging multiple spike trains (as captured by shuffled autocorrelation statistics, Joris 2003, or as might result from convergence of afferents from multiple fibers on the same neuron), attenuates some, but not all, effects of refractoriness (not shown).

In summary, this short paper suggests approaches for phenomenological modeling of the spike generation process, for the purpose of better understanding the properties of this spike-based code, to allow simulation of downstream processes that use this code, and possibly also to gain additional insight into the sound-to-spike transduction process itself.

## Notes

### Competing Interest Statement

The authors have declared no competing interest.

